# Predicting alcohol use from genome-wide polygenic scores, environmental factors, and their interactions in young adulthood

**DOI:** 10.1101/2020.07.05.188656

**Authors:** Radhika Kandaswamy, Andrea Allegrini, Alexandra F. Nancarrow, Sophie Nicole Cave, Robert Plomin, Sophie von Stumm

## Abstract

Alcohol use during emerging adulthood is associated with adverse life outcomes but its risk factors are not well known. Here, we predicted alcohol use in 3,153 young adults aged 22 years from (a) genome-wide polygenic scores (GPS) based on genome-wide association studies for the target phenotypes number of drinks per week and Alcohol Use Disorders Identification Test scores, (b) 30 environmental factors, and (c) their interactions (i.e., GxE effects). Data was collected from 1994 to 2018 as a part of the UK Twins Early Development Study. GPS accounted for up to 1.9% of the variance in alcohol use (i.e., Alcohol Use Disorders Identification Test score), while the 30 measures of environmental factors together accounted for 21.1%. The 30 GPS-environment interactions did not explain any additional variance and none of the interaction terms exceeded the significance threshold after correcting for multiple testing. Our findings suggest that GPS and environmental factors have primarily direct, additive effects rather than interacting systematically.

---

Alcohol use in emerging adulthood, including early initiation, frequent consumption, and drunkenness, is associated with adverse psychological and physical health outcomes (1-2). However, few environmental risk factors have been found that are causal predictors of alcohol use in emerging adulthood (3). Alcohol consumption is a complex behavioral trait influenced by many genetic variants called single-nucleotide polymorphisms (SNPs) that each have a very small effect size (4-5) and are detected in large genome-wide association (GWA) studies. The identified SNPs can then be aggregated into genome-wide polygenic scores (GPS) that capture a person’s genetic propensity for alcohol consumption.

Over the past few years, a number of GWA studies for alcohol consumption have been published with sample sizes ranging from ∼67,000 to nearly one million individuals with a vast range of target phenotypes, including the number of alcoholic drinks/units consumed per week (4, 6), alcohol intake in grams per day (5), maximum habitual alcohol intake (7), and alcohol use and dependence (i.e., Alcohol Use Disorder Identification Test (AUDIT) scores (8-9)). GPS based on these GWA studies predict alcohol use, with the variance accounted for ranging from 0.1% to 2.5% (4, 5, 10, 11). In general, the predictive power of a GPS depends on the GWA sample size, SNP heritability of the trait, selection thresholds for selecting markers for creating the score, etc., among many other factors (12).

GPS provide currently the best platform to directly and robustly test if a genetic predisposition for alcohol use develops differently across environmental contexts, a phenomenon known as gene-environment interactions (GxE; 13). Four previous studies tested GxE effects using GPS in the prediction of alcohol use, with mixed results. Two studies found no evidence for GxE: Using a GPS based on a GWA study of 67,000 individuals and their daily alcohol intake (5), no interactions were found between GPS and stress and life satisfaction in the prediction of alcohol use and alcohol-related problems in a sample of 6,705 Dutch adults (11). Similarly, no interaction was observed between peer drinking and GPS based on a GWA study of alcohol consumption in 112k UK Biobank participants (6) on alcohol use in a sample of 755 university students (14). However, two later studies reported significant GxE effects: Using GPS based on the publicly available summary statistics (i.e., excluding 23&Me data) from a GWA study of 535,000 individuals for the number of alcoholic drinks consumed per week (4), being in a romantic partnership was found to reduce the association between the genetic propensity for alcohol use and drinking frequency, intoxication frequency and alcohol dependence in 1201 young Finns (10). Also, the influence of GPS based on the GWA study mentioned above (6) on alcohol use per week was shown to be moderated by socioeconomic status (SES) in 6,567 Dutch adults (15). Alas, this finding could not be replicated with a GPS based on the GWAS & Sequencing Consortium of Alcohol and Nicotine use (GSCAN) study (4). Other studies that used ‘simpler’ GPS, which included fewer SNPs identified in smaller GWA study samples, suggested that genetic influences on alcohol use varied as a function of peer relations and parental knowledge (16-17). Overall, the current evidence for GxE in the prediction of alcohol use is inconclusive because studies employed different GWA studies to build GPS, including some with small sample sizes that afford low statistical power, and tested different environmental moderators, some of which could be argued to be heritable traits themselves (18), rendering direct comparisons of findings across studies impossible.

Here, we overcome previous studies’ limitations. For one, we used two large-scale, well-powered GWA studies to build GPS for alcohol use, including (1) the GSCAN study with an overall sample of 941,280 individuals that used the number of drinks consumed per week as target phenotype (4), and (2) the largest GWA study for AUDIT scores in 141,932 individuals (9). For the other, we considered many environments that were mostly assessed at age 22 and that are likely to influence the association between the genetic propensity for and actual alcohol consumption, rather than exploring only one or two environmental moderators. We note that few environments are truly exogenous to the individual (18), and many environmental measures show substantial genetic influences (19), with the possibility that environments and target phenotype (i.e. alcohol use) have shared their genetic aetiology (20).

We analyzed data from a large British cohort study, whose participants reported alcohol-related behaviors at age 22 and had data available on 30 environmental measures, which we categorized into four domains - home environment, adversity, lifestyles, and social and emotional learning competencies (SELC) – primarily to organize the results, rather than to propose a formal model for mapping out environments. SELC summarize traits of recognizing and managing emotions, setting and achieving positive goals, appreciating the perspectives of others, establishing and maintaining positive relationships, making responsible decisions, and handling interpersonal situations constructively (21). Our preregistered hypotheses included finding positive associations between the GPS and alcohol use, but we made no predictions about main effects of specific environments or the effect sizes associated with their GxE interactions due to the exploratory nature of our study.

## METHODS

### Sample

Data came from the Twins Early Development Study (TEDS; 22) that recruited families with twin births between 1994-1996 in England and Wales, who have been followed up in multiple waves until today. Although some attrition has occurred, TEDS remains largely representative of the UK population (22). Genotypes were processed using stringent quality control procedures followed by SNP imputation using the Haplotype Reference Consortium (release 1.1) reference panels. Following imputation, we excluded variants with minor allele frequency <0.5%, Hardy-Weinberg equilibrium *P-*values of <1 × 10^−5^. Full description of quality control and imputation procedures are provided in the Supplementary Methods. For this study we identified unrelated TEDS participants, for who genotype data were available at age 22 and who reported AUDIT-C scores (*N* = 3,390). We excluded 104 participants (3%), who reported never having had a drink containing alcohol, and 133 individuals (4%), who responded inconsistently to the AUDIT-C items. Our analysis sample consisted of a maximum of 3,153 individuals.

### Measures

#### Alcohol use

TEDS participants completed the 10 item AUDIT scale developed by the World Health Organization (WHO; 23) to identify hazardous and harmful alcohol use using paper, app- or web-based questionnaires at age 22. The first three items provide information pertaining to alcohol consumption and are often combined to create an alcohol subdomain score termed AUDIT-C. These items measure the frequency and quantity of typical alcohol drinking and the frequency of binge drinking. For the current analysis, the AUDIT-C score was created by aggregating the scores across all three items (see Supplementary Methods for details).

#### Environmental factors

Information on the environment measures was collected using paper, app-, or web-based questionnaires administered to TEDS participants at age 22 except SES, which was assessed at age 18 months (Supplementary Methods). In total, we included all available environmental factors in TEDS, which resulted in a comprehensive set of 30 quantitative, ordinal, and categorical measurements of environmental factors that represented four main domains: home environment, adversity, lifestyles, and social and emotional learning competencies (SELC) (Figure 1, Supplementary Table S1, Supplementary Methods). Home environment included (i) relationship with twin, (ii) CHAOS (confusion, hubbub and order scale), (iii) relationship status, (iv) whether on benefits, (v) education status, and (vi) parenthood. Adversity included (i) peer victimisation, (ii) life events, (iii) negative childhood experiences, and (iv) online bullying. Lifestyles included (i) conflict with the law, (ii) life satisfaction, (iii) healthy diet, (iv) BMI, (v) physical activity, (vi) involvement in sports, (vii) sleep quality, and (viii) online media use. SELC included social-emotional traits and behaviors, ranging from (i) self-control, (ii) risk-taking behavior, (iii) aggressive behavior, (iv) purpose in life, (v) volunteering, (vi) mood, (vii) peer pressure, (viii) ambition, (ix) general anxiety, (x) hassles to (xi) antisocial behavior. We note that this classification of the environmental variables is preliminary rather than confirmatory of a theoretical model. For a detailed description of these measures see Supplementary Methods. We used continuous measures of environments wherever possible to optimise the statistical power of our models.

**Figure 1.**
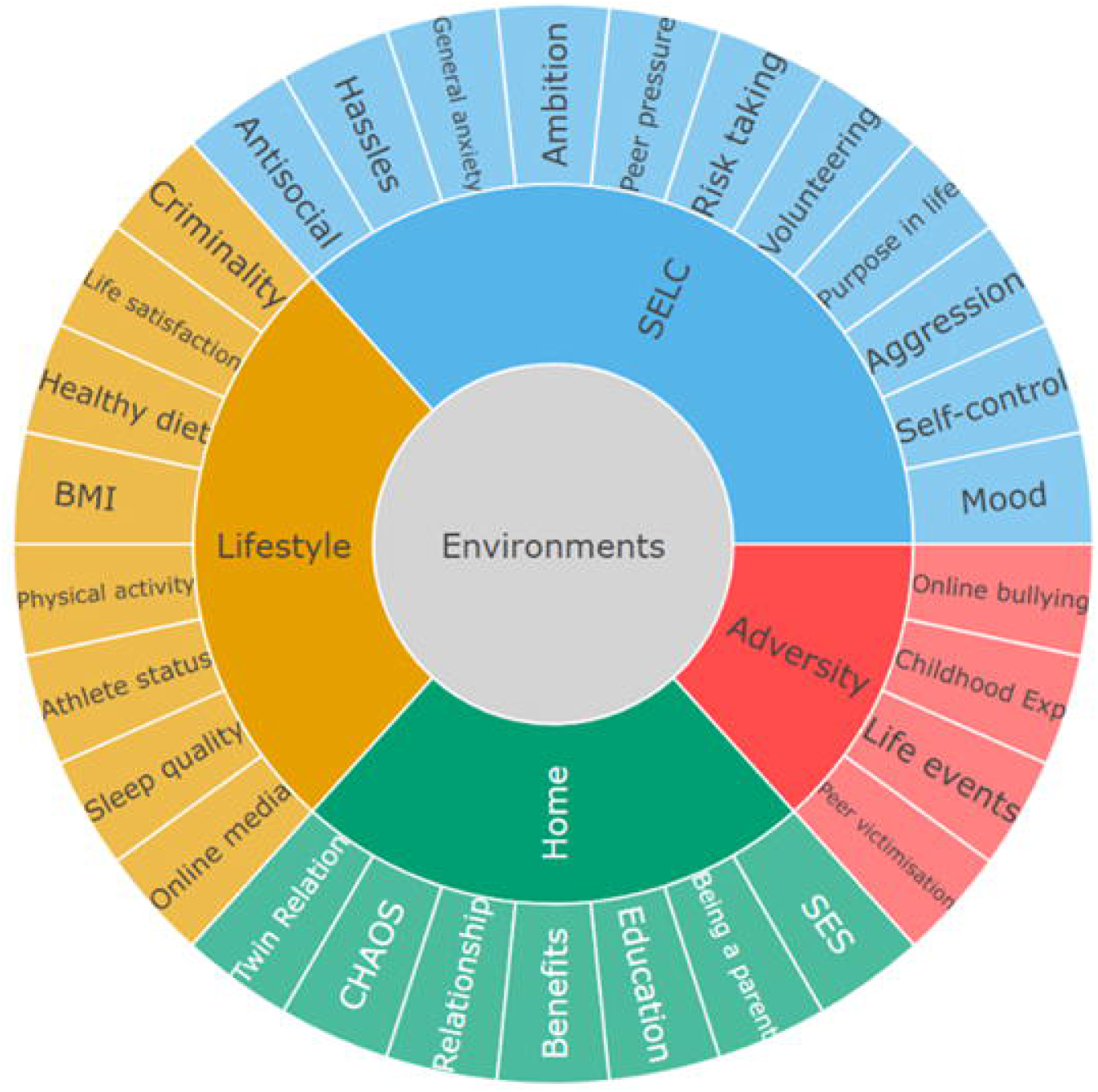
Summary of environment measures used in the present analyses of Twins Early Development Study participants. SELC = social and emotional learning competencies. SES = socioeconomic status. CHAOS = Confusion, Hubbub and Order Scale. BMI = body mass index.

### Genome-wide polygenic scores

The predictive power of a GPS depends on GWA study sample sizes, SNP heritability, and selection thresholds for selecting the markers for creating the score (12). For this reason we computed two GPS using PRSice-2 (24) and publicly available summary statistics from (i) the largest GSCAN study on “average number of alcoholic drinks consumed per week” that sampled ∼1 million individuals (after excluding 23andMe *N* = 537,349; 4), and (ii) the GWA study of AUDIT-C score (the same phenotype used in the current study) including UK Biobank research participants of European ancestry (after excluding 23andMe *N* = 121,604; 9). The GPS were generated as the sum of alleles associated with the phenotypic trait and weighted by their effect sizes reported in the discovery GWA studies. We performed clumping with r^2^ = 0.25 cut-off within a 250-kb window to remove SNPs in linkage disequilibrium and created eight scores for each discovery GWA study based on differing thresholds of discovery GWA study *P-*values (5e-08, 0.0001, 0.001, 0.01, 0.05, 0.1, 0.5, 1).

### Statistical analyses

First, we compared the predictive validity of each of the GPS created using the two GWA studies and differing *P*-value thresholds described in the section above for phenotypic alcohol use (AUDIT-C score) in our sample using individual linear regression models, controlling for gender, age, the first ten principal components (PCs; i.e., population stratification), and genotyping array. We then determined the GPS that explained the largest amount of variance in the target phenotype (based on Model R^2^, *P* < 0.05/16 tests = 0.003) and used that in all further analyses. Second, we tested the direct effects of each of the 30 environments on alcohol use in individual linear regressions models. Because our sample size is relatively small for the planned analyses, the *P*-value was Bonferroni-corrected to conservatively identify significant predictors (conventional *P* = 0.05/30 models = 0.002), instead of using a more lenient False Discovery Rate (FDR) method. We then performed a multiple linear regression including all 30 environments to predict alcohol use. Third, we individually tested moderation effects of each of the environments on the association between GPS and alcohol use (i.e., 30 models) by adding interaction terms to the regression models including the GPS and each environment, controlling again for gender, age, the first ten PCs, and genotyping array as covariates, as well as for all GPS x covariate and E x covariate interaction terms (25). Finally, we fitted a multiple regression model that included all interaction terms for the 30 environments and the GPS to assess their combined predictive power for alcohol use, controlling again for gender, age, the first ten PCs, and genotyping array as covariates, as well as for all GPS x covariate and E x covariate interaction terms. Predictor variables were scaled to mean 0 and unit variance prior to the analyses. All statistical analyses were conducted using R statistical software (26). All our analyses were preregistered at osf.io/ugk3n.

## RESULTS

### Genome-wide polygenic scores

Among the GPS generated using the GWA study for the number of drinks consumed per week (4) and for AUDIT-C scores (9) using eight selection *P*-value thresholds, 14 out of 16 exceeded the multiple testing significance threshold of *P* < 0.003 in the prediction of phenotypic alcohol use in our sample (Supplementary Figure S1, Supplementary Table S2). In general, the GPS based on both the GWA studies explained more variance in the phenotypic alcohol use as the selection *P*-value became less stringent. Specifically, GPS based on GSCAN study for the number of drinks consumed per week accounted for a maximum of 1.9% at the selection threshold of *P* < 0.1, while GPS based on the GWA study for AUDIT-C scores accounted for only 0.9% of the variance at best (Supplementary Figure S1, Supplementary Table S2). We therefore carried forward the best predictive GPS_*P* < 0.1_ based on GSCAN study for the number of drinks consumed per week (4) for all further analyses.

### Predicting alcohol use from environmental factors

Correlations between AUDIT-C and GSCAN GPS and the environmental factors (*P* < 0.01) are displayed in Supplementary Figure S2. When testing the individual, direct effects of the 30 environments on alcohol use, 14 exceeded the Bonferroni-corrected significance threshold (*Bonf P* < 0.002, Supplementary Table S3A-D, Figure 2A). From the domain of home environments (7 measures), being a parent, being on benefits and SES were associated with alcohol use, such that higher SES individuals consumed more alcohol and individuals with children or on benefits consumed less alcohol (Supplementary Table S3A). In the adversity domain (4 measures), only peer victimization was significantly associated with alcohol use, with individuals who experienced more victimization by peers consuming more alcohol (Supplementary Table S3B). From the lifestyles (8 measures), life satisfaction, physical activity, involvement in sports at a competitive level (i.e., athlete status) and online media use were associated with higher alcohol use (Supplementary Table S3C). Among the 11 factors in the SELC domain, lower self-control and higher risk taking, aggression, peer pressure, ambition and antisocial behavior were significantly associated with greater alcohol use (Supplementary Table S3D). The amount of variance explained by each of these environments ranged from 0.4% for ambition to 11.5% for risk-taking behavior (Supplementary Table S3A-D).

**Figure 2.**
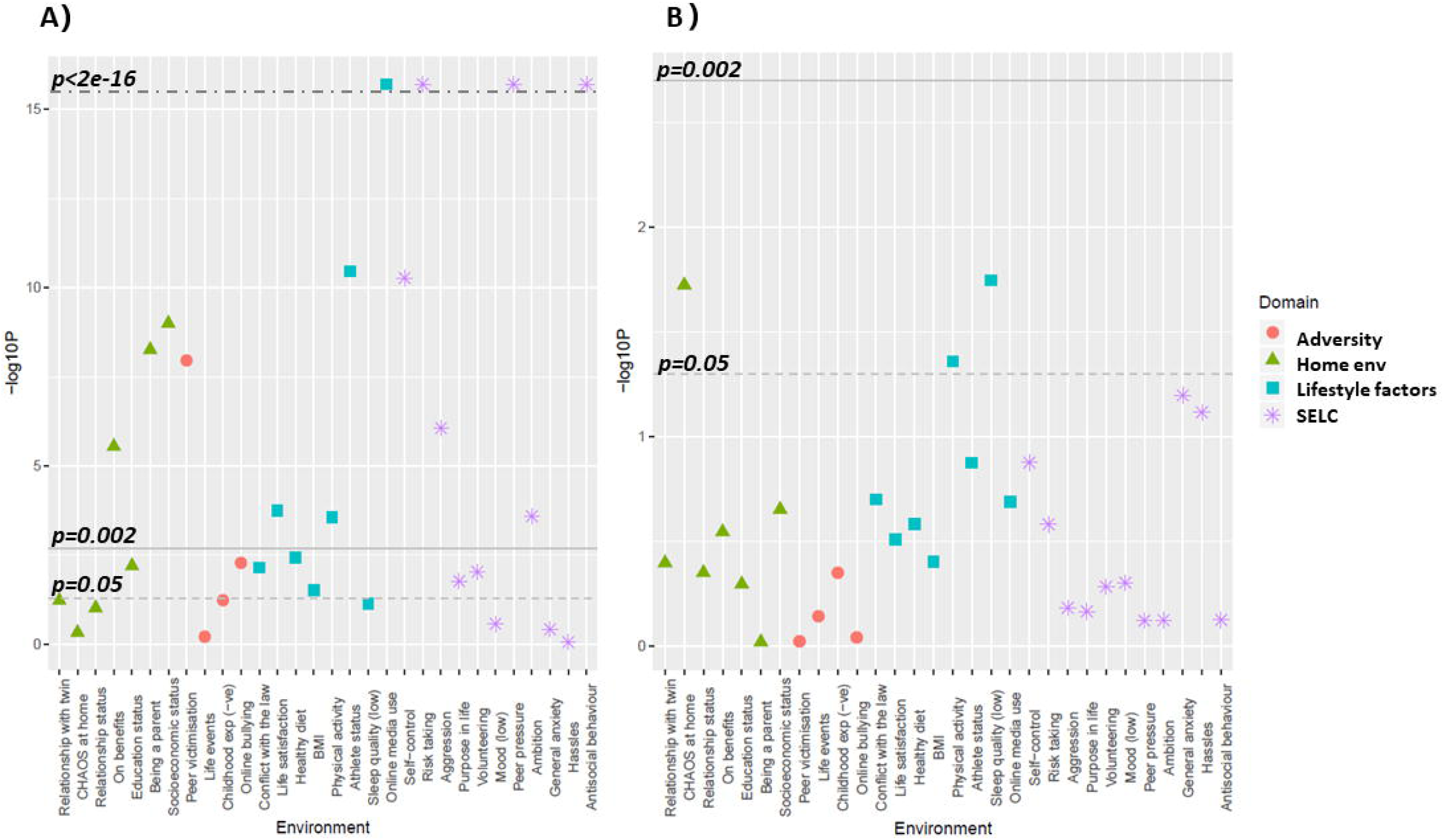
*P*-values for the associations between AUDIT-C scores and environments (A) and GxE interaction terms (B) from the univariate regression models for the present study of Twins Early Development Study participants. A) The *P*-values are based on the 30 individual regression models for each environment and its association with AUDIT-C scores, controlling for age, gender, PCs, and genotyping array. B) The *P*-values are based on the 30 individual regression models including the GPS, the environment, and their GxE interaction term, controlling for age, gender, PCs, genotyping array, G x covariates and E x covariates terms. In panels A and B, the grey dotted line refers to*P* < 0.05, while solid grey line refers to the Bonferroni-corrected *P* < 0.002, and dark grey dotted line refers to *P* < 2e-16, which is the lowest *P*-value that R reports. SELC, Social and emotional learning competencies.

Jointly, the 30 environments accounted for 21.1% of the variance in alcohol use. The environments that remained significant predictors in the multiple regression model included being a parent (home environment domain), online media use (lifestyles domain), and peer pressure, risk-taking and antisocial behavior (SELC domain; Supplementary Table S4). Akin to the univariate regression results, risk-taking emerged as the strongest predictor of alcohol use, independently accounting for 6% of the variance after adjusting for all other environmental effects. The GPS and all the 30 environments together accounted for 22.2% of the variance in alcohol use (Supplementary Table S5).

### Testing GxE in alcohol use

When testing the individual interaction effects of the GPS and the 30 environment factors on alcohol use, none exceeded the Bonferroni-corrected significance threshold (*Bonf P* < 0.002; (Supplementary Table S3A-D, Figure 2B). A multiple regression model with all the interaction terms between the GPS and each of the 30 environments together with the main effects of the GPS and the environments accounted for a total of 22.5% of the variance in alcohol use (Supplementary Table S6). However, the GxE interaction terms did not explain any additional variance in alcohol use beyond the confounding effects of G x covariates and E x covariates. None of the G x E interaction terms reached statistical significance after multiple testing correction (Supplementary Table S6).

## DISCUSSION

Our study is the first to offer a comprehensive analysis of the influence of 30 factors that map environments on the genetic contribution to alcohol use in emerging adulthood. We found that a GPS based on the GSCAN study for alcohol consumption (4) predicted alcohol use in our sample of young adults, but there was no evidence for moderation of this genetic effect by any of the environmental factors. The discrepancies between our current and previous findings (e.g., 10, 15) are likely to be due to the vast methodological differences between studies. In particular, previous studies in this area based their GPS on different GWA studies, assessed different phenotypes, and tested populations of different ages (10, 11, 14, 15).

Our findings contribute to a growing body of literature that struggles to identify GxE effects (27-30), because of three key challenges. First, studies that examine only one or two environmental moderators are likely to identify false positives, because of residual confounding that is unaccounted for (cf. 25). We partly overcome this challenge in the current research by comprehensively assessing the environment and testing 30 environmental moderators simultaneously. Second, testing GxE effects requires extremely large samples that afford sufficient statistical power to detect associations of very small effect size, which are likely to be true for GxE terms (25, 27). In the current study, we analyzed data from more than 3,000 individuals, which compares well with other GxE studies on alcohol use but is small relative to the populations that are typically assessed in GWA studies (31). Post-hoc power calculations using G*Power (32) suggested that our study had more than 80% power to detect two-way interactions with an effect size of f^2^=0.005 (15) but had less than 37% power to detect GxE when all the interaction terms are entered simultaneously with their direct effects. Because interaction models require much more power compared to tests of main effects, the modest predictive validity of the GPS for alcohol use in young adulthood suggests that the sample size required to detect GxE may be far greater than those currently available (28). Third, a principal caveat in GxE studies is the discrepancy in the assessment of genes and environment. GPS capture inherited DNA differences that are systematically coded in base pairs of single-nucleotide polymorphisms (SNPs) and remain unchanged from conception throughout life (33). By comparison, measures of the environment capture broad, often stochastic events that continuously change across time and contexts. Identifying interactions between two domains that are so asymmetrically assessed in research is inherently complicated, if not impossible.

Our results suggest that alcohol use in emerging adulthood is best predicted by a plethora of factors, including genetics with our GPS accounting for up to 1.9% of the variance in the AUDIT-C scores. Greater effect sizes were observed when modelling the environmental variables jointly: Together they accounted for ∼21% of the variance in alcohol use, with peer pressure, parenthood, online media use, risk-taking and antisocial behavior emerging as significant predictors. Risk-taking appeared as a particularly strong predictor, accounting independently for 6% of the variance in alcohol use. Risk-taking and alcohol use are genetically correlated (34), suggesting a common genetic origin. Future research will have to disentangle if both traits are causally related across development.

### Limitations

Our study has several limitations. First, the discovery GWA study for creating the GPS and the target sample were both composed mainly of European ancestry making the GxE findings from this study less generalizable to other ancestry populations. Second, TEDS like all longitudinal studies has suffered from attrition, although it still remains fairly representative of the population in England and Wales in terms of ethnicity and family SES (22). Third, we included 30 environmental factors in our analyses; yet, it is possible other environments that we did not assess play an important role in moderating the genetic contributions to alcohol use. Fourth, we Bonferroni-corrected our *P*-values for multiple testing, which might be an overly conservative and stringent approach, which avoids detecting false positives but limits observing small effect sizes. Finally, we did not explore sex-specific moderation by the environments on the genetic contributions to alcohol use behavior due to our lack of power to detect three-way interactions.

### Conclusion

We undertook here for the first time a hypothesis-free approach for identifying GxE interactions by investigating many environmental factors, instead of using one or two as moderators of the genetic predisposition to alcohol use. We confirmed previous findings that GPS predict alcohol use in young adults, with small effect sizes, and we found that not having a child, risk-taking, antisocial behavior, peer pressure, and use of online media increased the risk of alcohol use in emerging adulthood. However, we found no support for GxE effects on alcohol use in the current analyses. Future studies with larger sample sizes and more powerful polygenic scores, based on bigger GWA studies, that comprehensively assessed the environment are key to unravelling the complex interplay between the genes and environment in predicting alcohol use. Our results suggest that the effects of genetics and environments on alcohol use are primarily additive rather than due to interactive effects.

