## Supplementary Figure S1 for "Predicting alcohol use from genome-wide polygenic scores, environmental factors, and their interactions in young adulthood"

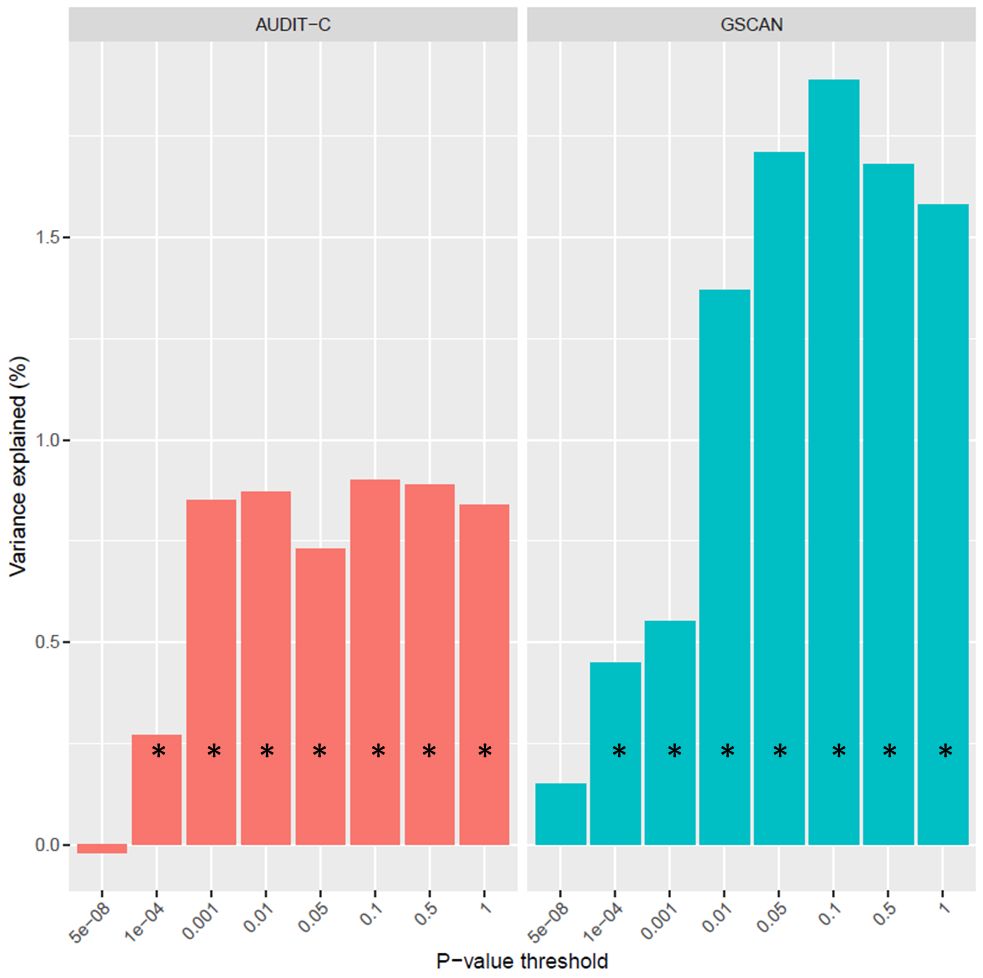

**Supplementary Figure S1** Predictive power of genome-wide polygenic scores based on AUDIT-C (Clarke *et al.,* 2017) (left panel) and GSCAN (*Liu et al., 2019*) (right panel) GWA studies. Vertical bars represent change in model R^2^ from baseline model (age, sex, first 10 principal components and genotyping array as covariates) to model including polygenic scores at various discovery GWA study *p*-value inclusion thresholds for AUDIT-C scores. **Bonferroni p* *< 0.003. GWA, genome-wide association study, GSCAN, Genome & Sequencing Consortium*.
