## Supplementary figures and images for "Predicting alcohol use from genome-wide polygenic scores, environmental factors, and their interactions in young adulthood"

### Supplementary Figure S2

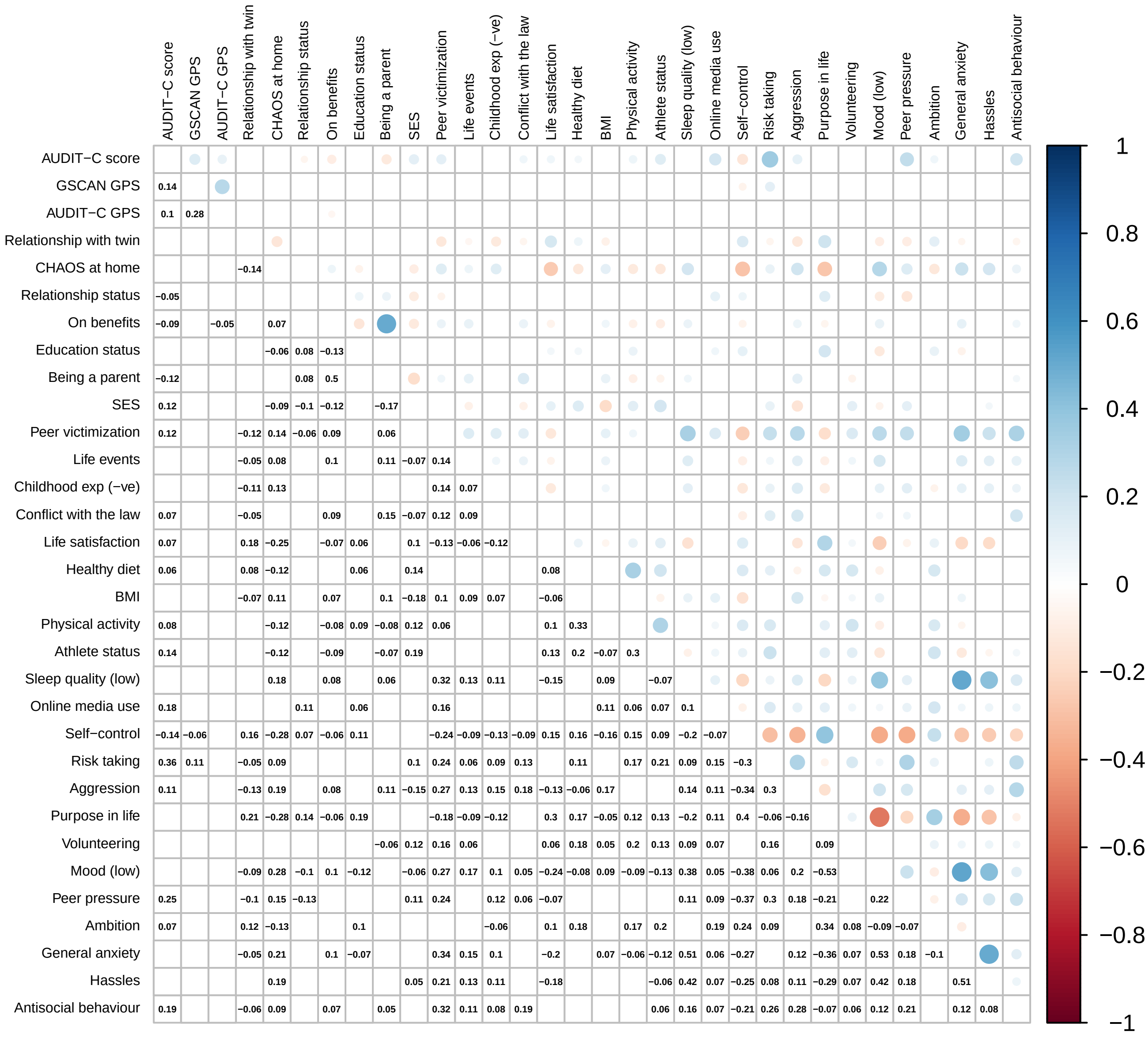
