## Supplementary Tables S1, S2, S3A, S3B, S3C, S3D, S4, S5, S6, S7 for "Predicting alcohol use from genome-wide polygenic scores, environmental factors, and their interactions in young adulthood"

**Supplementary Table S1. Descriptive statistics of environmental factors and AUDIT-C scores included in the study.**

| ***Domain*** | ***Environment*** | ***Category*** | **Mean / N** | **SD / %** |
| --- | --- | --- | --- | --- |
| **Home environment** | Relationship with twin | Continuous | 19.8 | 4.4 |
|  | CHAOS at home | Continuous | 3.7 | 2.2 |
|  | Relationship status | Categorical | 1360(0) vs1582(1) | 46% |
|  | On benefits | Categorical | 2788(0) vs149(1) | 95% |
|  | Education status (Unemployed vs All) | Categorical | 171(0) vs 2768(1) | 6% |
|  | Being a parent | Categorical | 2855 (0) vs 95 (1) | 97% |
|  | Socioeconomic status | Continuous | 0.3 | 1.0 |
| **Adversity** | Peer victimisation | Continuous | 3.7 | 5.0 |
|  | Affected by Life events | Categorical | 878 (0) vs 2269 (1) | 28% |
|  | Negative childhood experiences | Categorical | 428 (0) vs 2710 (1) | 14% |
|  | Online bullying | Categorical | 2109 vs 822 (1) | 0.4 |
| **Lifestyles** | Conflict with the law | Categorical | 2851 (0) vs 210 (1) | 93% |
|  | Life satisfaction | Continuous | 18.2 | 3.6 |
|  | Healthy diet | Continuous | 6.8 | 3.4 |
|  | BMI | Continuous | 23.4 | 4.5 |
|  | Physical activity | Continuous | 23.4 | 3.1 |
|  | Athlete status | Categorical | 0(757) vs 1(958) vs 2(1229) | 26% vs 33% vs 41% |
|  | Sleep quality | Continuous | 4.9 | 3.8 |
|  | Online media use | Continuous | 33.4 | 5.4 |
| **SELC** | Self-control | Continuous | 14.6 | 4.2 |
|  | Risk taking | Continuous | 6.5 | 3.4 |
|  | Aggression | Continuous | 19.8 | 6.1 |
|  | Purpose in life | Continuous | 17.3 | 4.0 |
|  | Volunteering | Continuous | 5.8 | 3.2 |
|  | Mood | Continuous | 4.4 | 4.2 |
|  | Peer pressure | Continuous | 13.8 | 5.0 |
|  | Ambition | Continuous | 13.8 | 3.6 |
|  | General anxiety | Continuous | 7.4 | 7.4 |
|  | Hassles | Continuous | 12.6 | 4.6 |
|  | Antisocial behaviour | Categorical | 2410 (0) vs 662 (1) | 78% |
| **Alcohol measure** | AUDIT-C score | Continuous | 5.7 | 2.8 |

*Note: Quantitative variables are described by average ± standard deviation. Ordinal variables are described by the number and percentage of individuals in each category. Education status was coded as a bivariate term, which contrasts being unemployed with being actively engaged in education, work, travel, or care, including studying, or working, apprenticeship or other employment training, gap year or travelling, or being a full-time parent.*

**Supplementary Table S2. Associations between all PRS fractions for alcohol consumption and AUDIT-C scores.**

| ***GWAS*** | ***p-value threshold*** | ***Beta*** | ***SE*** | ***p-value*** | ***CI_lower*** | ***CI_upper*** | ***∆R2 (%)*** |
| --- | --- | --- | --- | --- | --- | --- | --- |
| **AUDIT-C** | 5.00E-08 | 0.03 | 0.05 | 0.595 | -0.07 | 0.12 | -0.02 |
|  | 1.00E-04 | 0.15 | 0.05 | **0.002** | 0.06 | 0.25 | 0.27 |
|  | 1.00E-03 | 0.26 | 0.05 | **1.12E-07** | 0.17 | 0.36 | 0.85 |
|  | 0.01 | 0.27 | 0.05 | **7.74E-08** | 0.17 | 0.36 | 0.87 |
|  | 0.05 | 0.24 | 0.05 | **8.32E-07** | 0.15 | 0.34 | 0.73 |
|  | 0.1 | 0.27 | 0.05 | **4.48E-08** | 0.17 | 0.37 | 0.90 |
|  | 0.5 | 0.27 | 0.05 | **5.20E-08** | 0.17 | 0.37 | 0.89 |
|  | 1 | 0.26 | 0.05 | **1.14E-07** | 0.17 | 0.36 | 0.84 |
| **GSCAN** | 5.00E-08 | 0.12 | 0.05 | 0.015 | 0.02 | 0.22 | 0.15 |
|  | 1.00E-04 | 0.20 | 0.05 | **8.54E-05** | 0.10 | 0.29 | 0.45 |
|  | 1.00E-03 | 0.22 | 0.05 | **1.49E-05** | 0.12 | 0.31 | 0.55 |
|  | 0.01 | 0.33 | 0.05 | **1.91E-11** | 0.24 | 0.43 | 1.37 |
|  | 0.05 | 0.37 | 0.05 | **6.59E-14** | 0.28 | 0.47 | 1.71 |
|  | 0.1 | 0.39 | 0.05 | **3.27E-15** | 0.30 | 0.49 | 1.89 |
|  | 0.5 | 0.37 | 0.05 | **1.16E-13** | 0.27 | 0.47 | 1.68 |
|  | 1 | 0.36 | 0.05 | **5.85E-13** | 0.26 | 0.46 | 1.58 |

*Note: AUDIT-C GWAS (Clarke et al., 2017); GSCAN GWAS (Liu et al., 2019); Statistically significant p< 0.003 are presented in bold. All regression models were adjusted for sex, age, first 10 Principal components and genotyping array. Covariates only model presented in* ***Supplementary Table 7****; CI, confidence interval.*

**Supplementary Table 3A. Stepwise addition of variables to linear model predicting AUDIT-C scores. Effects of adding variables and interactions to the Covariates only model predicting variance in AUDIT-C scores, with environments from the Home environment domain.**

| ***Regression Models*** | | ***Beta*** | ***SE*** | ***p-value*** | ***Adj R2*** | ***Delta R2**** |
| --- | --- | --- | --- | --- | --- | --- |
| **Relationship with Twin** | | | | | | |
| Covariates only | |  | | | 0.0202 |  |
| Covariates + Relationship with twin | | 0.099 | 0.053 | 0.06 | 0.0211 | 0.001 |
| Covariates + GPS | | **0.409** | **0.051** | **1.69E-15** | 0.0410 | 0.021 |
| Covariates + Relationship with twin + GPS | E | 0.107 | 0.052 | 0.04 | 0.0420 | 0.022 |
|  | GPS | **0.411** | **0.051** | **1.23E-15** |  |  |
| Covariates + Relationship with twin + GPS + Relationship w Twin x GPS | E | 1.717 | 1.353 | 0.20 | 0.0401 | -9.9E-05 |
|  | GPS | 1.479 | 1.368 | 0.28 |  |  |
|  | GxE | -0.045 | 0.053 | 0.40 |  |  |
| **CHAOS at home** | | | | | | |
| Covariates only | |  | | | 0.0194 |  |
| Covariates + CHAOS at home | | -0.037 | 0.051 | 0.47 | 0.0192 | -1.6E-04 |
| Covariates + GPS | | **0.400** | **0.051** | **3.41E-15** | 0.0393 | 0.020 |
| Covariates + CHAOS at home + GPS | E | -0.039 | 0.051 | 0.44 | 0.0391 | 0.020 |
|  | GPS | **0.400** | **0.051** | **3.34E-15** |  |  |
| Covariates + CHAOS at home + GPS + CHAOS x GPS | E | 2.427 | 1.355 | 0.07 | 0.0429 | 0.001 |
|  | GPS | 1.695 | 1.358 | 0.21 |  |  |
|  | GxE | -0.126 | 0.053 | 0.02 |  |  |
| **Relationship status** | | | | | | |
| Covariates only | |  | | | 0.0198 |  |
| Covariates + Relationship status | | -0.172 | 0.104 | 0.10 | 0.0204 | 0.001 |
| Covariates + GPS | | **0.403** | **0.051** | **3.71E-15** | 0.0399 | 0.020 |
| Covariates + Relationship status + GPS | E | -0.166 | 0.102 | 0.11 | 0.0405 | 0.021 |
|  | GPS | **0.402** | **0.051** | **4.06E-15** |  |  |
| Covariates + Relationship status + GPS + Relationship status x GPS | E | -0.438 | 2.648 | 0.87 | 0.0411 | -1.4E-04 |
|  | GPS | 1.634 | 1.363 | 0.23 |  |  |
|  | GxE | -0.079 | 0.103 | 0.45 |  |  |
| **On Benefits** | | | | | | |
| Covariates only | |  | | | 0.0197 |  |
| Covariates + On benefits | | **-1.096** | **0.234** | **2.8E-06** | 0.0267 | 0.007 |
| Covariates + GPS | | **0.412** | **0.051** | **1.1E-15** | 0.0407 | 0.021 |
| Covariates + On benefits + GPS | E | **-1.108** | **0.231** | **1.7E-06** | 0.0479 | 0.028 |
|  | GPS | **0.413** | **0.051** | **6.6E-16** |  |  |
| Covariates + On benefits + GPS + On benefits x GPS | E | 4.798 | 6.073 | 0.43 | 0.0484 | 4.6E-05 |
|  | GPS | 1.560 | 1.363 | 0.25 |  |  |
|  | GxE | -0.254 | 0.238 | 0.29 |  |  |
| **Education status** | | | | | | |
| Covariates only | |  | | | 0.0195 |  |
| Covariates + Education | | 0.599 | 0.219 | 0.01 | 0.0216 | 0.002 |
| Covariates + GPS | | **0.412** | **0.051** | **8.7E-16** | 0.0406 | 0.021 |
| Covariates + Education + GPS | E | 0.606 | 0.217 | 0.01 | 0.0429 | 0.023 |
|  | GPS | **0.413** | **0.051** | **7.3E-16** |  |  |
| Covariates + Education + GPS + Education x GPS | E | -0.133 | 6.332 | 0.98 | 0.0404 | -1.9E-04 |
|  | GPS | 1.256 | 1.376 | 0.36 |  |  |
|  | GxE | 0.148 | 0.223 | 0.51 |  |  |
| **Being a parent** | | | | | | |
| Covariates only | |  | | | 0.0193 |  |
| Covariates + Parent | | **-1.699** | **0.290** | **5.4E-09** | 0.0303 | 0.011 |
| Covariates + GPS | | **0.409** | **0.051** | **1.3E-15** | 0.0401 | 0.021 |
| Covariates + Parent + GPS | E | **-1.767** | **0.287** | **8.7E-10** | 0.0520 | 0.033 |
|  | GPS | **0.418** | **0.051** | **2.1E-16** |  |  |
| Covariates + Parent + GPS + Parent x GPS | E | -2.043 | 8.104 | 0.80 | 0.0515 | -3.3E-04 |
|  | GPS | 1.389 | 1.368 | 0.31 |  |  |
|  | GxE | -0.014 | 0.274 | 0.96 |  |  |
| **Socioeconomic status (SES)** | | | | | | |
| Covariates only | |  | | | 0.0200 |  |
| Covariates + SES | | **0.317** | **0.052** | **1.0E-09** | 0.0318 | 0.012 |
| Covariates + GPS | | **0.408** | **0.050** | **6.7E-16** | 0.0408 | 0.021 |
| Covariates + SES + GPS | E | **0.311** | **0.051** | **1.4E-09** | 0.0522 | 0.032 |
|  | GPS | **0.404** | **0.050** | **9.2E-16** |  |  |
| Covariates + SES + GPS + SES x GPS | E | 2.813 | 1.339 | 0.04 | 0.0513 | 1.6E-04 |
|  | GPS | 1.010 | 1.347 | 0.45 |  |  |
|  | GxE | 0.062 | 0.051 | 0.22 |  |  |

*Note: Significant terms (p<0.002) are in bold. Interactions include all main effects, covariates and covariate interaction terms. E, Environment; GPS, Genome-wide polygenic scores.*

**Delta Rsquared for the E, GPS and E + GPS models is the increase in variance explained compared to the covariates only model. Delta Rsquared for the GxE interaction term is the increase in variance compared to a model with main effects of GPS, Environment, the interaction terms including the G x Cov and E x Cov terms but without the GxE interaction term.*

**Supplementary Table 3B. Stepwise addition of variables to linear model predicting AUDIT-C scores. Effects of adding variables and interactions to the Covariates only model predicting variance in AUDIT-C scores, with environments from the Adversity domain.**

| ***Regression Models*** | | ***Beta*** | ***SE*** | ***p-value*** | ***Adj R2*** | ***Delta R2**** |
| --- | --- | --- | --- | --- | --- | --- |
| **Peer victimisation** | | | | | | |
| Covariates only | | | | | 0.0202 |  |
| Covariates + Peer victimisation | | **0.288** | **0.050** | **1.11E-08** | 0.0303 | 0.010 |
| Covariates + GPS | | **0.391** | **0.050** | **7.54E-15** | 0.0391 | 0.019 |
| Covariates + Peer victimisation + GPS | E | **0.283** | **0.050** | **1.51E-08** | 0.0489 | 0.029 |
|  | GPS | **0.387** | **0.050** | **1.03E-14** |  |  |
| Covariates + Peer victimisation + GPS + Peer victimisation x GPS | E | 2.264 | 1.302 | 0.08 | 0.0499 | -3.13E-04 |
|  | GPS | 0.926 | 1.318 | 0.48 |  |  |
|  | GxE | -0.003 | 0.050 | 0.96 |  |  |
| **Affected by Life events** | | | | | | |
| Covariates only | | | | | 0.0190 |  |
| Covariates + Life events | | 0.053 | 0.111 | 0.63 | 0.0187 | -0.0002 |
| Covariates + GPS | | **0.391** | **0.050** | **3.90E-15** | 0.0378 | 0.019 |
| Covariates + Life events + GPS | E | 0.049 | 0.110 | 0.65 | 0.0376 | 0.019 |
|  | GPS | **0.391** | **0.050** | **4.01E-15** |  |  |
| Covariates + Life events + GPS + Life events x GPS | E | 2.450 | 2.824 | 0.39 | 0.0363 | -2.71E-04 |
|  | GPS | 1.388 | 1.321 | 0.29 |  |  |
|  | GxE | -0.040 | 0.113 | 0.72 |  |  |
| **Negative childhood experiences** | | | | | | |
| Covariates only | | | | | 0.0190 |  |
| Covariates + Neg Childhood Exp. | | 0.273 | 0.145 | 0.06 | 0.0198 | 0.001 |
| Covariates + GPS | | **0.394** | **0.050** | **2.65E-15** | 0.0381 | 0.019 |
| Covariates + Neg Childhood Exp. + GPS | E | 0.259 | 0.144 | 0.07 | 0.0388 | 0.020 |
|  | GPS | **0.393** | **0.050** | **3.10E-15** |  |  |
| Covariates + Neg Childhood Exp. + GPS + Neg Childhood Exp. x GPS | E | 4.152 | 3.756 | 0.27 | 0.0335 | -1.35E-04 |
|  | GPS | 1.374 | 1.327 | 0.30 |  |  |
|  | GxE | 0.115 | 0.153 | 0.45 |  |  |
| **Online bullying** | | | | | | |
| Covariates only | | | | | 0.0198 |  |
| Covariates + Online bullying | | 0.318 | 0.114 | 0.01 | 0.0221 | 0.002 |
| Covariates + GPS | | **0.408** | **0.051** | **1.89E-15** | 0.0405 | 0.021 |
| Covariates + Online bullying + GPS | E | 0.282 | 0.113 | 0.01 | 0.0422 | 0.022 |
|  | GPS | **0.403** | **0.051** | **4.19E-15** |  |  |
| Covariates + Online bullying + GPS + Online bullying x GPS | E | 4.205 | 2.917 | 0.15 | 0.0459 | -3.27E-04 |
|  | GPS | 1.704 | 1.360 | 0.21 |  |  |
|  | GxE | 0.012 | 0.112 | 0.92 |  |  |

*Note: Significant terms (p<0.002) are in bold. Interactions include all main effects, covariates and covariate interaction terms. E, Environment; GPS, Genome-wide polygenic scores.*

**Delta Rsquared for the E, GPS and E + GPS models is the increase in variance explained compared to the covariates only model. Delta Rsquared for the GxE interaction term is the increase in variance compared to a model with main effects of GPS, Environment, the interaction terms including the G x Cov and E x Cov terms but without the GxE interaction term.*

**Supplementary Table S3C. Stepwise addition of variables to linear model predicting AUDIT-C scores. Effects of adding variables and interactions to the Covariates only model predicting variance in AUDIT-C scores, with environments from the Lifestyles domain.**

| ***Regression Models*** | | ***Beta*** | ***SE*** | ***p-value*** | ***Adj R2*** | ***Delta R2**** |
| --- | --- | --- | --- | --- | --- | --- |
| **Conflict with the law** | | | | | | |
| Covariates only | |  |  |  | 0.0195 |  |
| Covariates + Conflict with law | | 0.541 | 0.201 | 0.01 | 0.0216 | 0.002 |
| Covariates + GPS | | **0.389** | **0.050** | **1.01E-14** | 0.0383 | 0.019 |
| Covariates + Conflict with law + GPS | E | 0.526 | 0.199 | 0.01 | 0.0402 | 0.021 |
|  | GPS | **0.388** | **0.050** | **1.17E-14** |  |  |
| Covariates + Conflict with law + GPS + Conflict with law x GPS | E | 3.415 | 4.988 | 0.49 | 0.0385 | 2.05E-04 |
|  | GPS | 0.986 | 1.329 | 0.46 |  |  |
|  | GxE | -0.261 | 0.204 | 0.20 |  |  |
| **Life satisfaction** | | | | | | |
| Covariates only | |  |  |  | 0.0191 |  |
| Covariates + Life satisfaction | | **0.191** | **0.051** | **1.77E-04** | 0.0234 | 0.004 |
| Covariates + GPS | | **0.409** | **0.051** | **1.27E-15** | 0.0399 | 0.021 |
| Covariates + Life satisfaction + GPS | E | **0.183** | **0.050** | **2.82E-04** | 0.0439 | 0.025 |
|  | GPS | **0.406** | **0.051** | **2.00E-15** |  |  |
| Covariates + Life satisfaction + GPS + Life satisfaction x GPS | E | 0.708 | 1.363 | 0.60 | 0.0419 | 1.04E-05 |
|  | GPS | 1.578 | 1.362 | 0.25 |  |  |
|  | GxE | -0.051 | 0.050 | 0.31 |  |  |
| **Healthy diet** | | | | | | |
| Covariates only | |  |  |  | 0.0206 |  |
| Covariates + Diet | | **0.156** | **0.054** | **3.64E-03** | 0.0234 | 0.003 |
| Covariates + GPS | | **0.401** | **0.053** | **8.04E-14** | 0.0407 | 0.020 |
| Covariates + Diet + GPS | E | **0.164** | **0.053** | **2.07E-03** | 0.0438 | 0.023 |
|  | GPS | **0.404** | **0.053** | **4.81E-14** |  |  |
| Covariates + Diet + GPS + Diet x GPS | E | 0.330 | 1.404 | 0.81 | 0.0435 | 9.44E-05 |
|  | GPS | 1.918 | 1.429 | 0.18 |  |  |
|  | GxE | 0.062 | 0.055 | 0.26 |  |  |
| **BMI** | | | | | | |
| Covariates only | |  |  |  | 0.0223 |  |
| Covariates + BMI | | -0.113 | 0.052 | 0.03 | 0.0236 | 0.001 |
| Covariates + GPS | | **0.400** | **0.052** | **1.85E-14** | 0.0424 | 0.020 |
| Covariates + BMI + GPS | E | -0.118 | 0.052 | 0.02 | 0.0439 | 0.022 |
|  | GPS | **0.401** | **0.052** | **1.46E-14** |  |  |
| Covariates + BMI + GPS + BMI x GPS | E | -1.704 | 1.406 | 0.23 | 0.0439 | -9.83E-05 |
|  | GPS | 1.346 | 1.396 | 0.34 |  |  |
|  | GxE | 0.044 | 0.052 | 0.40 |  |  |
| **Physical activity** | | | | | | |
| Covariates only | |  |  |  | 0.0194 |  |
| Covariates + Physical activity | | **0.187** | **0.051** | **2.68E-04** | 0.0235 | 0.004 |
| Covariates + GPS | | **0.412** | **0.051** | **9.07E-16** | 0.0405 | 0.021 |
| Covariates + Physical activity + GPS | E | **0.178** | **0.051** | **4.68E-04** | 0.0442 | 0.025 |
|  | GPS | **0.408** | **0.051** | **1.55E-15** |  |  |
| Covariates + Physical activity + GPS + Physical activity x GPS | E | -0.011 | 1.309 | 0.99 | 0.0425 | 0.001 |
|  | GPS | 1.326 | 1.363 | 0.33 |  |  |
|  | GxE | 0.104 | 0.052 | 0.04 |  |  |
| **Athlete status** | | | | | | |
| Covariates only | |  |  |  | 0.0195 |  |
| Covariates + Athlete status | | **0.425** | **0.064** | **3.5E-11** | 0.0345 | 0.015 |
| Covariates + GPS | | **0.413** | **0.051** | **6.8E-16** | 0.0407 | 0.021 |
| Covariates + Athlete status + GPS | E | **0.420** | **0.063** | **3.6E-11** | 0.0551 | 0.036 |
|  | GPS | **0.410** | **0.051** | **7.1E-16** |  |  |
| Covariates + Athlete status + GPS + Athlete status x GPS | E | -2.009 | 1.626 | 0.22 | 0.0580 | 0.001 |
|  | GPS | 1.837 | 1.351 | 0.17 |  |  |
|  | GxE | -0.096 | 0.064 | 0.13 |  |  |
| **Sleep** | | | | | | |
| Covariates only | |  |  |  | 0.0201 |  |
| Covariates + Sleep | | 0.092 | 0.051 | 0.07 | 0.0208 | 0.001 |
| Covariates + GPS | | **0.385** | **0.050** | **1.49E-14** | 0.0383 | 0.018 |
| Covariates + Sleep + GPS | E | 0.093 | 0.050 | 0.07 | 0.0391 | 0.019 |
|  | GPS | **0.385** | **0.050** | **1.39E-14** |  |  |
| Covariates + Sleep + GPS + Sleep x GPS | E | 1.085 | 1.311 | 0.41 | 0.0414 | 0.001 |
|  | GPS | 1.477 | 1.326 | 0.27 |  |  |
|  | GxE | 0.124 | 0.052 | 0.02 |  |  |
| **Online media use** | | | | | | |
| Covariates only | |  |  |  | 0.0200 |  |
| Covariates + Online media use | | **0.518** | **0.050** | **2.39E-24** | 0.0539 | 0.034 |
| Covariates + GPS | | **0.410** | **0.051** | **1.26E-15** | 0.0410 | 0.021 |
| Covariates + Online media use + GPS | E | **0.505** | **0.050** | **1.13E-23** | 0.0732 | 0.053 |
|  | GPS | **0.394** | **0.050** | **5.96E-15** |  |  |
| Covariates + Online media use + GPS + Online media use x GPS | E | 2.140 | 1.294 | 0.10 | 0.0720 | 1.96E-04 |
|  | GPS | 1.356 | 1.343 | 0.31 |  |  |
|  | GxE | -0.066 | 0.052 | 0.20 |  |  |

*Note: Significant terms (p<0.002) are in bold. Interactions include all main effects, covariates and covariate interaction terms. E, Environment; GPS, Genome-wide polygenic scores.*

**Delta Rsquared for the E, GPS and E + GPS models is the increase in variance explained compared to the covariates only model. Delta Rsquared for the GxE interaction term is the increase in variance compared to a model with main effects of GPS, Environment, the interaction terms including the G x Cov and E x Cov terms but without the GxE interaction term.*

**Supplementary Table S3D. Stepwise addition of variables to linear model predicting AUDIT-C scores. Effects of adding variables and interactions to the Covariates only model predicting variance in AUDIT-C scores, with environments from the Social and emotional learning competencies (SELC) domain.**

| ***Regression Models*** | | ***Beta*** | ***SE*** | ***p-value*** | ***Adj R2*** | ***Delta R2*** |
| --- | --- | --- | --- | --- | --- | --- |
| **Self-control** | | | | | | |
| Covariates only | |  |  |  | 0.0175 |  |
| Covariates + Self-control | | **-0.344** | **0.052** | **5.4E-11** | 0.0320 | 0.014 |
| Covariates + GPS | | **0.412** | **0.052** | **2.6E-15** | 0.0386 | 0.021 |
| Covariates + Self-control + GPS | E | **-0.319** | **0.052** | **8.7E-10** | 0.0510 | 0.033 |
|  | GPS | **0.391** | **0.052** | **4.2E-14** |  |  |
| Covariates + Self-control + GPS + Self-control x GPS | E | -3.165 | 1.341 | 0.02 | 0.0505 | 4.24E-04 |
|  | GPS | 0.866 | 1.380 | 0.53 |  |  |
|  | GxE | 0.080 | 0.053 | 0.13 |  |  |
| **Risk-taking** | | | | | | |
| Covariates only | |  |  |  | 0.0176 |  |
| Covariates + Risk-taking | | **0.965** | **0.050** | **1.1E-78** | 0.1323 | 0.115 |
| Covariates + GPS | | **0.408** | **0.052** | **4.7E-15** | 0.0383 | 0.021 |
| Covariates + Risk-taking + GPS | E | **0.930** | **0.050** | **1.2E-73** | 0.1437 | 0.126 |
|  | GPS | **0.305** | **0.049** | **6.1E-10** |  |  |
| Covariates + Risk-taking + GPS + Risk-taking x GPS | E | 0.801 | 1.285 | 0.53 | 0.1462 | 7.73E-05 |
|  | GPS | 1.288 | 1.311 | 0.33 |  |  |
|  | GxE | 0.058 | 0.052 | 0.26 |  |  |
| **Aggression** | | | | | | |
| Covariates only | |  |  |  | 0.0176 |  |
| Covariates + Aggression | | **0.259** | **0.052** | **8.8E-07** | 0.0256 | 0.008 |
| Covariates + GPS | | **0.408** | **0.052** | **4.7E-15** | 0.0383 | 0.021 |
| Covariates + Aggression + GPS | E | **0.249** | **0.052** | **1.7E-06** | 0.0457 | 0.028 |
|  | GPS | **0.402** | **0.052** | **9.3E-15** |  |  |
| Covariates + Aggression + GPS + Aggression x GPS | E | 0.984 | 1.334 | 0.46 | 0.0463 | -2.74E-04 |
|  | GPS | 0.799 | 1.380 | 0.56 |  |  |
|  | GxE | 0.024 | 0.053 | 0.66 |  |  |
| **Purpose in life** | | | | | | |
| Covariates only | |  |  |  | 0.0167 |  |
| Covariates + Purpose in Life | | 0.123 | 0.051 | 0.02 | 0.0183 | 0.002 |
| Covariates + GPS | | **0.403** | **0.051** | **3.5E-15** | 0.0369 | 0.020 |
| Covariates + Purpose in Life + GPS | E | 0.124 | 0.051 | 0.01 | 0.0385 | 0.022 |
|  | GPS | **0.403** | **0.051** | **3.1E-15** |  |  |
| Covariates + Purpose in Life + GPS + Purpose in Life x GPS | E | -0.841 | 1.311 | 0.52 | 0.0388 | -2.76E-04 |
|  | GPS | 1.574 | 1.361 | 0.25 |  |  |
|  | GxE | -0.021 | 0.052 | 0.68 |  |  |
| **Volunteering** | | | | | | |
| Covariates only | |  |  |  | 0.0193 |  |
| Covariates + Volunteering | | 0.134 | 0.051 | 0.01 | 0.0213 | 0.002 |
| Covariates + GPS | | **0.416** | **0.051** | **5.6E-16** | 0.0409 | 0.022 |
| Covariates + Volunteering + GPS | E | 0.125 | 0.051 | 0.01 | 0.0425 | 0.023 |
|  | GPS | **0.414** | **0.051** | **8.2E-16** |  |  |
| Covariates + Volunteering + GPS + Volunteering x GPS | E | 2.582 | 1.354 | 0.06 | 0.0405 | -1.96E-04 |
|  | GPS | 1.381 | 1.363 | 0.31 |  |  |
|  | GxE | -0.034 | 0.052 | 0.52 |  |  |
| **Mood** | | | | | | |
| Covariates only | |  |  |  | 0.0194 |  |
| Covariates + Mood | | -0.058 | 0.052 | 0.27 | 0.0195 | 7.98E-05 |
| Covariates + GPS | | **0.415** | **0.051** | **6.6E-16** | 0.0408 | 0.021 |
| Covariates + Mood + GPS | E | -0.055 | 0.051 | 0.28 | 0.0408 | 0.021 |
|  | GPS | **0.415** | **0.051** | **7.0E-16** |  |  |
| Covariates + Mood + GPS + Mood x GPS | E | 2.659 | 1.322 | 0.04 | 0.0418 | -1.81E-04 |
|  | GPS | 1.273 | 1.363 | 0.35 |  |  |
|  | GxE | 0.035 | 0.053 | 0.50 |  |  |
| **Peer Pressure** | | | | | | |
| Covariates only | |  |  |  | 0.0194 |  |
| Covariates + Peer pressure | | **0.647** | **0.051** | **1.9E-36** | 0.0712 | 0.052 |
| Covariates + GPS | | **0.415** | **0.051** | **6.6E-16** | 0.0408 | 0.021 |
| Covariates + Peer pressure + GPS | E | **0.631** | **0.050** | **2.1E-35** | 0.0900 | 0.071 |
|  | GPS | **0.390** | **0.050** | **7.0E-15** |  |  |
| Covariates + Peer pressure + GPS + Peer pressure^a^ x GPS | E | 1.353 | 1.309 | 0.30 | 0.0857 | -2.87E-04 |
|  | GPS | 1.681 | 1.335 | 0.21 |  |  |
|  | GxE | -0.016 | 0.051 | 0.76 |  |  |
| **Ambition** | | | | | | |
| Covariates only | |  |  |  | 0.0190 |  |
| Covariates + Ambition | | **0.182** | **0.050** | **2.5E-04** | 0.0228 | 0.004 |
| Covariates + GPS | | **0.391** | **0.050** | **3.9E-15** | 0.0378 | 0.019 |
| Covariates + Ambition + GPS | E | **0.185** | **0.049** | **1.7E-04** | 0.0418 | 0.023 |
|  | GPS | **0.393** | **0.049** | **2.7E-15** |  |  |
| Covariates + Ambition + GPS + Ambition x GPS | E | -2.460 | 1.286 | 0.06 | 0.0388 | -2.79E-04 |
|  | GPS | 1.411 | 1.320 | 0.29 |  |  |
|  | GxE | -0.016 | 0.050 | 0.76 |  |  |
| **General Anxiety** | | | | | | |
| Covariates only | |  |  |  | 0.0206 |  |
| Covariates + General Anxiety | | -0.045 | 0.051 | 0.38 | 0.0205 | -7.14E-05 |
| Covariates + GPS | | **0.391** | **0.050** | **5.2E-15** | 0.0395 | 0.019 |
| Covariates + General Anxiety + GPS | E | -0.038 | 0.050 | 0.45 | 0.0393 | 0.019 |
|  | GPS | **0.391** | **0.050** | **5.8E-15** |  |  |
| Covariates + General Anxiety + GPS + General Anxiety x GPS | E | 1.027 | 1.290 | 0.43 | 0.0395 | 7.66E-04 |
|  | GPS | 1.293 | 1.325 | 0.33 |  |  |
|  | GxE | 0.095 | 0.051 | 0.06 |  |  |
| **Hassles** | | | | | | |
| Covariates only | |  |  |  | 0.0190 |  |
| Covariates + Hassles | | -0.008 | 0.050 | 0.88 | 0.0187 | -3.06E-04 |
| Covariates + GPS | | **0.391** | **0.050** | **3.9E-15** | 0.0378 | 0.019 |
| Covariates + Hassles + GPS | E | -0.017 | 0.050 | 0.73 | 0.0375 | 0.019 |
|  | GPS | **0.392** | **0.050** | **3.8E-15** |  |  |
| Covariates + Hassles + GPS + Hassles x GPS | E | 0.460 | 1.320 | 0.73 | 0.0359 | 0.001 |
|  | GPS | 1.359 | 1.329 | 0.31 |  |  |
|  | GxE | 0.091 | 0.052 | 0.08 |  |  |
| **Antisocial behaviour** | | | | | | |
| Covariates only | |  |  |  | 0.0202 |  |
| Covariates + Antisocial behavior | | **1.186** | **0.122** | **3.5E-22** | 0.0495 | 0.029 |
| Covariates + GPS | | **0.392** | **0.050** | **6.4E-15** | 0.0391 | 0.019 |
| Covariates + Antisocial behavior + GPS | E | **1.146** | **0.121** | **4.0E-21** | 0.0664 | 0.046 |
|  | GPS | **0.371** | **0.049** | **7.4E-14** |  |  |
| Covariates + Antisocial behavior + GPS + Antisocial behavior x GPS | E | 4.830 | 3.034 | 0.11 | 0.0639 | -2.78E-04 |
|  | GPS | 1.130 | 1.308 | 0.39 |  |  |
|  | GxE | -0.039 | 0.125 | 0.75 |  |  |

*Note: Significant terms (p<0.002) are in bold. Interactions include all main effects, covariates and covariate interaction terms. E, Environment; GPS, Genome-wide polygenic scores.*

*^a^A single item assessing peer pressure for getting drunk in parties did not show any interaction effect as well (data not shown).*

**Delta Rsquared for the E, GPS and E + GPS models is the increase in variance explained compared to the covariates only model. Delta Rsquared for the GxE interaction term is the increase in variance compared to a model with main effects of GPS, Environment, the interaction terms including the G x Cov and E x Cov terms but without the GxE interaction term.*

**Supplementary Table S4. Associations between all environments and AUDIT-C scores using multiple linear regression model.**

| ***Domain*** | ***Environment*** | ***Beta*** | ***SE*** | ***p-value*** | ***Adj Rsquared*** |
| --- | --- | --- | --- | --- | --- |
| **Home env** | Relationship with twin | 0.123 | 0.058 | 0.033 | 0.229 |
|  | CHAOS at home | -0.016 | 0.059 | 0.787 |  |
|  | Relationship status | -0.128 | 0.110 | 0.244 |  |
|  | On benefits | 0.047 | 0.320 | 0.883 |  |
|  | Education status | 0.258 | 0.232 | 0.268 |  |
|  | Being a parent | **-1.341** | **0.426** | **0.002** |  |
|  | Socioeconomic status | 0.122 | 0.059 | 0.040 |  |
| **Adversity** | Peer victimisation | 0.098 | 0.068 | 0.150 |  |
|  | Life events | -0.068 | 0.119 | 0.569 |  |
|  | Negative childhood exp | 0.077 | 0.159 | 0.627 |  |
|  | Online bullying | -0.020 | 0.132 | 0.880 |  |
| **Lifestyles** | Conflict with the law | 0.454 | 0.230 | 0.048 |  |
|  | Life satisfaction | 0.129 | 0.057 | 0.025 |  |
|  | Healthy diet | -0.028 | 0.058 | 0.629 |  |
|  | BMI | -0.022 | 0.013 | 0.079 |  |
|  | Physical activity | -0.114 | 0.061 | 0.061 |  |
|  | Athlete status | 0.165 | 0.074 | 0.026 |  |
|  | Sleep quality (low) | 0.030 | 0.068 | 0.657 |  |
|  | Online media use | **0.344** | **0.057** | **2.49E-09** |  |
| **SELC** | Self-control | -0.160 | 0.070 | 0.023 |  |
|  | Risk taking | **0.803** | **0.064** | **2.00E-16** |  |
|  | Aggression | -0.042 | 0.063 | 0.506 |  |
|  | Purpose in life | 0.053 | 0.072 | 0.459 |  |
|  | Volunteering | -0.148 | 0.058 | 0.010 |  |
|  | Mood (low) | -0.160 | 0.074 | 0.030 |  |
|  | Peer pressure | **0.327** | **0.060** | **6.60E-08** |  |
|  | Ambition | 0.118 | 0.060 | 0.050 |  |
|  | General anxiety | 0.024 | 0.076 | 0.748 |  |
|  | Hassles | -0.185 | 0.068 | 0.006 |  |
|  | Antisocial behaviour | **0.641** | **0.145** | **1.08E-05** |  |

*Note: Baseline Model (Cov only) Rsquared=0.01873, p-value = 2.84e-10; Statistically significant results (p<0.002) are presented in bold; SELC, Social and emotional learning competencies*

**Supplementary Table S5. Associations between GPS, all environments and AUDIT-C scores using multiple linear regression model.**

| ***Domain*** | ***Environment*** | ***Beta*** | ***SE*** | ***p-value*** | **Adj. *Rsquared*** |
| --- | --- | --- | --- | --- | --- |
| **Home env** | Relationship with twin | 0.136 | 0.058 | 0.018 | 0.240 |
|  | CHAOS at home | -0.012 | 0.059 | 0.842 |  |
|  | Relationship status | -0.114 | 0.109 | 0.299 |  |
|  | On benefits | 0.072 | 0.318 | 0.821 |  |
|  | Education status | 0.266 | 0.231 | 0.249 |  |
|  | Being a parent | **-1.395** | **0.423** | **0.001** |  |
|  | Socioeconomic status | 0.123 | 0.059 | 0.038 |  |
| **Adversity** | Peer victimisation | 0.105 | 0.068 | 0.122 |  |
|  | Life events | -0.055 | 0.118 | 0.640 |  |
|  | Negative childhood exp | 0.068 | 0.158 | 0.669 |  |
|  | Online bullying | -0.049 | 0.131 | 0.708 |  |
| **Lifestyles** | Conflict with the law | 0.458 | 0.228 | 0.045 |  |
|  | Life satisfaction | 0.121 | 0.057 | 0.034 |  |
|  | Healthy diet | -0.013 | 0.058 | 0.829 |  |
|  | BMI | -0.023 | 0.013 | 0.071 |  |
|  | Physical activity | -0.129 | 0.060 | 0.032 |  |
|  | Athlete status | 0.162 | 0.074 | 0.028 |  |
|  | Sleep quality (low) | 0.032 | 0.067 | 0.635 |  |
|  | Online media use | **0.340** | **0.057** | **2.85E-09** |  |
| **SELC** | Self-control | -0.144 | 0.070 | 0.039 |  |
|  | Risk taking | **0.771** | **0.063** | **< 2e-16** |  |
|  | Aggression | -0.035 | 0.063 | 0.575 |  |
|  | Purpose in life | 0.051 | 0.072 | 0.472 |  |
|  | Volunteering | -0.149 | 0.057 | 0.010 |  |
|  | Mood (low) | -0.150 | 0.073 | 0.040 |  |
|  | Peer pressure | 0.330 | 0.060 | **3.97E-08** |  |
|  | Ambition | 0.123 | 0.060 | 0.039 |  |
|  | General anxiety | 0.035 | 0.075 | 0.642 |  |
|  | Hassles | -0.196 | 0.067 | 0.003 |  |
|  | Antisocial behaviour | **0.616** | **0.144** | **2.09E-05** |  |
| **GPS** | | **0.291** | **0.053** | **4.41E-08** |  |

*Note: Baseline Model (Cov only) Rsquared=0.01873, p-value = 2.84e-10; SELC, Social and emotional learning competencies; GPS, Genome-wide polygenic scores*

**Supplementary Table S6. Associations between all GxE interaction terms and AUDIT-C scores in a multiple linear regression model.**

| ***Domain*** | | ***Environment*** | ***Beta*** | ***SE*** | ***p-value*** | **Adj. *Rsquared*** |
| --- | --- | --- | --- | --- | --- | --- |
| **GPS x** | **Home env** | Relationship with twin | 0.070 | 0.066 | 0.288 | 0.281 |
|  |  | CHAOS at home | -0.085 | 0.070 | 0.225 |  |
|  |  | Relationship status | -0.021 | 0.120 | 0.864 |  |
|  |  | On benefits | -0.181 | 0.384 | 0.637 |  |
|  |  | Education status | 0.257 | 0.245 | 0.294 |  |
|  |  | Being a parent | 0.181 | 0.521 | 0.728 |  |
|  |  | Socioeconomic status | 0.005 | 0.066 | 0.933 |  |
|  | **Adversity** | Peer victimisation | -0.108 | 0.077 | 0.161 |  |
|  |  | Life events | -0.029 | 0.134 | 0.831 |  |
|  |  | Negative childhood exp | -0.156 | 0.188 | 0.406 |  |
|  |  | Online bullying | 0.079 | 0.145 | 0.585 |  |
|  | **Lifestyle** | Conflict with the law | -0.660 | 0.265 | 0.013 |  |
|  |  | Life satisfaction | 0.002 | 0.063 | 0.974 |  |
|  |  | Healthy diet | 0.025 | 0.068 | 0.712 |  |
|  |  | BMI | 0.073 | 0.064 | 0.260 |  |
|  |  | Physical activity | 0.010 | 0.069 | 0.889 |  |
|  |  | Athlete status | -0.088 | 0.082 | 0.285 |  |
|  |  | Sleep quality (low) | 0.060 | 0.082 | 0.465 |  |
|  |  | Online media use | -0.103 | 0.064 | 0.109 |  |
|  | **SELC** | Self-control | -0.021 | 0.079 | 0.786 |  |
|  |  | Risk taking | 0.158 | 0.076 | 0.036 |  |
|  |  | Aggression | 0.013 | 0.071 | 0.860 |  |
|  |  | Purpose in life | 0.113 | 0.081 | 0.165 |  |
|  |  | Volunteering | -0.047 | 0.066 | 0.480 |  |
|  |  | Mood (low) | 0.108 | 0.084 | 0.200 |  |
|  |  | Peer pressure | -0.059 | 0.068 | 0.386 |  |
|  |  | Ambition | -0.016 | 0.068 | 0.810 |  |
|  |  | General anxiety | 0.157 | 0.090 | 0.083 |  |
|  |  | Hassles | -0.001 | 0.075 | 0.990 |  |
|  |  | Antisocial behaviour | -0.073 | 0.166 | 0.661 |  |

*Note: The interaction terms included Gxcovariate and Excovariate terms along with the GxE terms. A Model with GPS, all Environments, Covariates and GxE but no GxCov and Excov terms Rsquared=0.2435, p-value = 2.2e-16; SELC, Social and emotional learning competencies; GPS, Genome-wide polygenic scores*

**Supplementary Table S7. Results for covariates only baseline model in the prediction of AUDIT-C scores.**

| ***Covariate*** | ***Beta*** | ***SE*** | ***p-value*** |
| --- | --- | --- | --- |
| Sex | **0.78** | **0.10** | **6.00E-14** |
| Age | -0.11 | 0.06 | 0.05 |
| PC1 | -10.31 | 5.27 | 0.05 |
| PC2 | 8.29 | 5.31 | 0.12 |
| PC3 | -2.62 | 5.39 | 0.63 |
| PC4 | 4.43 | 5.39 | 0.41 |
| PC5 | 5.26 | 5.46 | 0.33 |
| PC6 | -4.65 | 5.48 | 0.40 |
| PC7 | 2.53 | 5.42 | 0.64 |
| PC8 | -2.15 | 5.47 | 0.69 |
| PC9 | 10.10 | 5.39 | 0.06 |
| PC10 | 6.83 | 5.42 | 0.21 |
| Genotyping Array | -0.03 | 0.10 | 0.78 |

*Note: Model adj. Rsquared: 0.01873, p = 2.84E-10*
