## Supplementary Methods for "Predicting alcohol use from genome-wide polygenic scores, environmental factors, and their interactions in young adulthood"

**Alcohol measures**

TEDS participants completed the 10 item AUDIT scale developed by World Health Organization (WHO) to identify hazardous and harmful alcohol use (1) using paper, app- or web-based questionnaires at age 22. The first three items of this tool measure the frequency and quantity of usual drinking and the frequency of binge drinking. For the current analysis, the AUDIT-C score was created by aggregating the scores from items 1-3. Data on alcohol measures and genotype was available for 3,390 unrelated individuals. 97% of the participants reported ever having had a drink containing alcohol, we excluded those who stated not to have consumed alcohol (*N*=104) from our analyses. Individuals with strong inconsistencies in responses to the three alcohol related items (Total *N*=133) were also removed from the analysis.

*Recoding of AUDIT Item 2 (alcohol consumption on a typical day):*

In the TEDS 22 questionnaire, this item was composed of 4 sub-items – measuring number of drinks of (a) Standard glass of wine (b) Pint of lager/beer/cider (c) Alcopop and (d) Single shot of spirit. These items were coded on a scale of 0-7 representing (0, 1-2, 3-5, 6-10, 11-15, 16-20, 21-25, 26 or more drinks). We used the midpoint of the range of the drinks and multiplied it by the number of units (2 for wine and beer, 1 for alcopop and spirit) to get the total number of units of each type of alcohol. We added these units from the four sub-items to obtain the total number of units of alcohol an individual consumed on a typical day. Individuals with total values >5 standard deviations from the mean (≥37 units) were set to missing (*N*=29). Final sample consisted of 3153 individuals with information on genotypes and AUDIT-C outcome.

**Description of the environment and lifestyle factors included in the study**

Information on questionnaires with the TEDS environment variables can be accessed at https://www.teds.ac.uk/datadictionary/studies/measures/21yr_measures.htm

***A. Home environment***

1. *Twin relationship:* The quality of the sibling relationship assessed using five items on a 5 point scale from strongly disagree to strongly agree: *“I enjoy my relationship with my twin”, “My twin and I have a lot of fun together”, “I like to spend time with my twin”, “My twin and I do a lot of things together”,* and *“My twin talks to me about personal problems”.* A mean score was created and multiplied by the total number of items to represent this variable, with higher values indicating better twin relationships.
2. *Chaos at home:* CHAOS (Confusion, Hubbub and Order Scale) at home. A home environment scale assessing the household disorganization using six items (2). Data were collected from twins, who rated statements on a 3-point scale from not true to very true. The six items are: *“It’s chaotic in the house”, “You can’t hear yourself think in the house”, “There is regular routine in the morning”, “Everyone in the house is usually able to stay on top of things”, “There is usually a television turned on somewhere in the house” and “The atmosphere in the house is calm”.* A mean score was created and multiplied by the total number of items to represent this variable, with higher values indicating higher disorder or chaos at home.
3. *Relationship status:* An item to assess the current relationship status was recoded (0 vs 1) such that categories - married, in exclusive relationship, and living with partner were merged into one (1) and all others into not being currently in a relationship (0), including single, dating non-exclusively, widowed, separated, and divorced.
4. *On benefits or not*: Assessed using a single item - whether the individual is currently receiving benefits (Yes/No).
5. *Education status:* Assessed using a single item “Which of the following best describes what you are currently doing? (tick one only)” Responses included: “Studying”, “Working”, “Apprenticeship or other employment training”, “Gap year/ travelling”, “Unemployed”, and “Full time parent”. The responses were recoded with unemployed as 0 and all other responses coded as 1.
6. *Being a parent or not:* Assessed using a single item asking if the twin had a child (Yes/No).
7. *Socio economic status:* SES at first contact (mean age = 1.5 years) was calculated as a composite of mother and father qualification levels ranging from 1 = ‘no qualifications’ to 8 = ‘postgraduate qualification’, mother and father employment status, and mothers’ age at first birth.

***B. Adversity***

1. *Peer victimisation:* Assessed using 16 items on self-reported victimisation experiences since age 16 by peers based on the Multidimensional Peer-Victimization Scale (3). It contains three factors – Verbal and physical victimisation, and social manipulation. A mean score was created and multiplied by the total number of items to represent this variable, with higher values indicating higher peer victimisation.
2. *Life events:* Assessed life events experienced since age 16 using 11 items on a scale of 0-4 if the event had happened or not (coded as 0) and Yes, affected me a lot (4). Items included *“You became homeless”, “You or your partner became pregnant or had a baby”, “You lost your job or got into serious financial problems “, “You were divorced or separated”, “You were admitted to the hospital or became seriously ill”, “You were in trouble with the law”, “You were the victim of a serious crime”, “Someone close to you died”, “You attempted suicide”, “You or your partner had an abortion”, and “Your parents divorced*. A mean score was created from all the available responses to these items and recoded such that all individuals with a score > 0 were grouped into one category indicating two groups with and without life-events.
3. *Childhood experiences:* Negative childhood experiences were assessed using 8 items and a mean score was created for the analysis. Example items included are: *“When you were a child, how often did an adult in your family shout at you?”, “When you were a child, how often did an adult in your family shout at you?”, …. “When you were a child, how often did an adult hit you so hard it left you with bruises or marks?”* The items were coded from 0 – 4, from never to very often. A mean score was created from all the available responses and recoded such that all individuals with a score > 0 were grouped into one category indicating two groups with and without negative childhood experiences.
4. *Online bullying:* The questionnaire included four items for assessing experiences of bullying in childhood such as *“How often has someone sent you a nasty text (excluding family or partner)?”, “How often has someone written nasty things to you using instant messenger, such as Facebook Messenger, Whatsapp, Snapchat (excluding family or partner)?”* Items were scored from 0-2, for not at all, once and more than once. A mean of the responses to the four items was created and this item was recoded such that all individuals with a score > 0 were grouped into one category indicating two groups with and without online bullying experience for the analysis.

***C. Lifestyle factors***

5. *Crime or conflict with the law:* We used three items of the total six items assessing criminality at age 22 to get a mean score. The items are *“Have you ever been cautioned by the police?”, “Have you ever been arrested?” and “Have you ever been sentenced to prison?”*. A mean of the responses to the four items was created and this item was recoded such that all individuals with a score > 0 were grouped into one category indicating two groups with and without conflict with the law for the analysis
6. *Life satisfaction:* This measure assesses an individual's feeling of community within their neighbourhood using 5 items based on the CLAS Life Satisfaction Scale – Community (4). A mean score was generated and multiplied by the number of total items in the scale for the analysis.
7. *Health behaviours/Healthy diet:* Questionnaire was designed to assess healthy eating and lifestyle behaviours using 12 items. Out of these only 4 items corresponding to healthy eating behaviour were used *(a) eating 3 portions of whole grain products, (b) 5 portions of fruits and vegetable, (c) 3-4 portions of milk and dairy foods or dairy alternatives and (d) 2 portions of protein-rich foods in one day*. The items were coded from 0-4 for rarely, 1-2 times per week, 3-4 times per week, 5-6 times per week and every day. A mean score was created from the items and multiplied by the total number of items. There was no clear separation of the individuals based on eating healthy vs not in our data, hence we decided to use mean total score rather than two categories as planned in our preregistration document.
8. *BMI: Body Mass Index (BMI),* derived from self-reported height and weight was used. BMI = weight (kg)/height(m)^2^.
9. *Physical activity*: A brief questionnaire of 3 items assessing physical activity levels during a typical week, specifically assessing mild, moderate and strenuous exercise. Each kind of exercise was coded from 1-5 from 0-15 mins (coded as 1), 16-60 mins (coded as 2), 61-120 mins (coded as 3), 121-180 mins (coded as 4) and 181+ mins (coded as 5). We recoded the responses to moderate and strenuous exercise such that it was continuous with 0-15mins of moderate exercise coded as 6 and continuing the same pattern till strenuous exercise 3+ hours as 15. Finally, a mean score of all the scores for mild, moderate and strenuous level of exercise was created and multiplied by the total number of items to indicate level of physical activity.
10. *Involvement in sports:* A single item assessing engagement of the twin in any sport at a competitive level. Responses were recoded (0-2) such that no participation was 0, merge participated in sport at a social or non-competitive level and competed within organised individual sport events as 1, and all other responses for competed in school to international level as 2.
11. *Sleep quality*: Nine items assessing sleep quality based on the Pittsburgh Sleep Quality Index (5). Example items: *“During the past month, how often have you had trouble sleeping because you (1) Woke up in the middle of the night or early morning? (2) Had to get up to use the bathroom? ….(8) Had bad dreams? (9) Had pain?”.* Items were coded on a scale of 0-3 from Not during the past month to Three or more times per week. We created a mean score from all the items and multiplied it with the total number of items in the scale for our analysis.
12. *Online media use:* 11 items assessing online behaviour and online media usage on a 6 point scale from 0-5 (Never, several times a year, Several times a month, several times a week, several times a day and several times an hour). Example items: *How often do you … (1) send, receive and read e-mails? (2) send and receive text messages or check for text messages? (3) make and receive calls on your mobile phone? … (6) watch video clips? (7) play games by yourself, with other people in the same room, or with other people online? …..(11) Comment or click ‘like’ on postings, status updates, photos, etc.* A mean score from the responses to these items was created and multiplied by the total number of items in the scale to represent online behaviour.

***D. Social and emotional learning competencies:***

13. *Self-control:* A mean score was created by averaging up the responses to six items measuring self-control, adapted from the Brief Self-Control measure (6) and shortened from 13 to 6 items . For eg., *“I am good at resisting temptation, I have a hard time breaking bad habits, I am lazy etc*.*”* The mean score was multiplied by the total number of items in the scale to get a composite score. Higher scores represented higher self-control.
14. *Risk-taking index:* A mean score was created by averaging the responses for six items measuring recreational risks, health risks, career risks, financial risks, safety risks, and social risks and multiplied by the total number of items in the scale (7).
15. *Aggression:* A measure consisting of 8 items based on the Brief Aggression Questionnaire (BAQ; 8) assessing verbal and physical aggression. A mean score was created and multiplied by the total number of items in the scale to represent overall aggressive behaviour.
16. *Purpose in life:* Assessed using 5 items shortened from 20 items with responses on a scale of 1-5 (9). Items included are such as *“I feel my personal existence is (1) Utterly meaningless, without purpose to (5) Purposeful and meaningful”, “As I view the world in relation to my life, the world* …with responses from on a scale of 1-5 with Completely confuses me and Fits meaningfully with my life at each end.” A mean score was created from the responses to the items and multiplied by the total number of items in the scale and used in our analysis.
17. *Volunteering:* This measure is composed of 5 items looking at money spent by individuals on charity or money given to beggars, and if they provided unpaid help to charity or people during the last 12 months. Example items: *“How often have you given money to charity?”, “How often have you sponsored a friend who was raising money for charity?”, “How often have you given unpaid help to other people?”.* The responses to all these items were averaged and multiplied by the total number of items in the scale to indicate level of volunteering*.*
18. *Mood:* Assessed using eight items based on the Short Mood and Feeling Questionnaire (SMFQ) (Shortened from 13 to 8 items assessing mood in the last two weeks (10). Example items are *“I felt miserable or unhappy, I felt so tired I just sat around and did nothing, I was very restless, I thought I could never be as good as other people etc!* with responses from (0) Not true to (2) very true. A mean score from the responses to all the items was created and multiplied by the total number of items to have a continuous scale representing mood in the last two weeks.
19. *Peer pressure:* This measure assessed the perception of peer pressure in different scenarios using 7 items such as *“I give in to peer pressure easily, At times, I have broken rules because others have urged me to, At times I have done dangerous or foolish things because others dared me, I have felt pressured to get drunk at parties etc”* with responses from (0) Strongly disagree to (5) Strongly agree (11). A mean score was created and multiplied by the total number of items for the analysis. Item specifically assessing alcohol-related peer pressure – “I have felt pressured to get drunk at parties” was used separately to see if it interacted with the GPS of Alcohol consumption.
20. *Ambition:* This measure contains 5 items such as *“I aim to be the best in the world at what I do, I am ambitious, Achieving something of lasting importance is the highest goal in life, I think achievement is overrated and I am driven to succeed”.* The items were coded from 0-4 for Not at all like me, Not much like me, Somewhat like me, Mostly like me and Very much like me. A mean score was created from the items and multiplied by the total number of items. Looking at the distribution of the scores, we found it difficult to separate individuals with and without ambitions in our data based on an arbitrary cutoff, hence we decided to use mean total score rather than two categories as planned in our preregistration document.
21. *General anxiety:* Assessed using ten items assessing general anxiety (12) during the past 7 days, we generated an average based on all the responses to the items and multiplied it by the total number of items in the scale. Higher scores represented higher anxious behaviour.
22. *Hassles:* Hassles are irritants or things that annoy or bother individuals. Twins were asked seven questions about things that can be hassles in day-to-day life such as inner concerns, finances, time management, work, environment, family, and health on a scale of 0-4 from none or not applicable to A great deal (13). A mean of the responses available from these items was created and multiplied by the total number of items as a measure of hassles in day to day life.
23. *Antisocial behaviour:* A total of 16 items measuring antisocial behaviour during the last one year ranging from passive behaviour such as *“bought something that you knew or suspected was stolen” to active behaviour such as “How often have you been rowdy or rude in a public place, so that people complained or you got into trouble?”* (14). We created a mean score from the available item responses and multiplied by the total number of items to generate the composite score for our analysis.

**Detailed Genotyping and QC**

DNA for 8,743 individuals (including 3,722 dizygotic co-twin individuals) was extracted from saliva and buccal cheek swab samples and hybridized to HumanOmniExpressExome-8v1.2 genotyping arrays at the Institute of Psychiatry, Psychology and Neuroscience Genomics & Biomarker Core Facility. The raw image data from the array were normalized, pre-processed, and filtered in GenomeStudio according to Illumina Exome Chip SOP v1.4. (<http://confluence.brc.iop.kcl.ac.uk:8090/display/PUB/Production+Version%3A+Illumina+Exome+Chip+SOP+v1.4>). In addition, prior to genotype calling, 919 multi-mapping SNPs and 501 samples with callrate <0.95 were removed. The ZCALL program was used to augment the genotype calling for samples and SNPs that passed the initial QC.

DNA from 3,747 samples was extracted from buccal cheek swabs and genotyped at Affymetrix, Santa Clara, California, USA. From this sample, 3,665 samples were successfully hybridized to AffymetrixGeneChip 6.0 SNP genotyping arrays (<http://www.affymetrix.com/support/technical/datasheets/genomewide_snp6_datasheet.pdf>) using experimental protocols recommended by the manufacturer (Affymetrix Inc., Santa Clara, CA). The raw image data from the arrays were normalized and pre-processed at the Wellcome Trust Sanger Institute, Hinxton, UK for genotyping as part of the Wellcome Trust Case Control Consortium 2 (<https://www.wtccc.org.uk/ccc2/>) according to the manufacturer’s guidelines (http://www.affymetrix.com/support/downloads/manuals/genomewidesnp6_manual.pdf). Genotypes for the Affymetrix arrays were called using CHIAMO (https://mathgen.stats.ox.ac.uk/genetics_software/chiamo/chiamo.html).

After initial quality control and genotype calling, the same quality control was performed on the samples genotyped on the Illumina and Affymetrix platforms separately using PLINK (15-16), R (17), BCFtools (18), and EIGENSOFT (19-20).

Samples were removed from subsequent analyses on the basis of call rate (<0.98), suspected non-European ancestry, heterozygosity, and relatedness other than dizygotic twin status. SNPs were excluded if the minor allele frequency was smaller than 0.5%, if more than 2% of genotype data were missing, or if the Hardy Weinberg *p*-value was lower than 10^-5^. Non-autosomal markers and indels were removed. Association between SNP and the platform, batch, plate or well on which samples were genotyped was calculated; SNPs with an effect *p*-value < 10^-4^ were excluded. A total sample of 10,346 samples (including 3,320 dizygotic twin pairs and 7,026 unrelated individuals), with 7,289 individuals and 559,772 SNPs genotyped on Illumina and 3,057 individuals and 635,269 SNPs genotyped on Affymetrix remained after quality control.

Genotypes from the two platforms were separately phased using EAGLE2 (21), and imputed into the Haplotype Reference Consortium (release 1.1) using the Positional Burrows-Wheeler Transform method (22) through the Sanger Imputation Service (23). Prior to merging, we excluded variants with info <0.75 and removed non-overlapping SNPs between platforms. After merging, we tested for minor allele frequency differences between platforms and removed SNPs with an effect p-value < 10^-4^, and Hardy Weinberg p-value > 10^-5^. Using these criteria, 7,363,646 genotyped and well-imputed SNPs were retained for the analyses.

To generate principal components to be used as covariates, we performed principal component analysis on a subset of 39,353 common (MAF > 5%), perfectly imputed (info = 1) autosomal SNPs, after stringent pruning to remove markers in linkage disequilibrium (r^2^ > 0.1) and excluding high linkage disequilibrium genomic regions so as to ensure that only genome-wide effects were detected*.*
